# Protection from Omicron and other VOCs by Bivalent S-Trimer COVID-19 Vaccine

**DOI:** 10.1101/2022.05.03.490428

**Authors:** Danmei Su, Xinglin Li, Xueqin Huang, Cui He, Cheng Zeng, Qiang Wang, Wenchang Qin, Zhongquan Mu, Donna Ambrosino, George Siber, Ralf Clemens, Joshua G. Liang, Peng Liang, Nick Jackson, Rong Xu

## Abstract

The Omicron variant of SARS-COV-2 (GISAID GRA clade [B.1.1.529, BA.1 and BA.2]) is now the single dominant Variant of Concern (VOC). The high number of mutations in the Omicron Spike (S) protein promotes humoral immunological escape. Although a third homologous boost with S, derived from the ancestral strain, was able to increase neutralizing antibody titers and breadth including to Omicron, the magnitude of virus neutralization could benefit from further optimization. Moreover, combining SARS-COV-2 strains as additional valences may address the current antigenicity range occupied by VOCs.

Using Trimer-Tag™ platform we have previously demonstrated phase 3 efficacy and safety of a prototypic vaccine SCB-2019 in the SPECTRA trial and have submitted applications for licensure. Here, we successfully generated a bivalent vaccine candidate including both Ancestor and Omicron variant S-proteins. Preclinical studies demonstrate this SARS-CoV-2 bivalent S-Trimer subunit vaccine elicits high titers of neutralizing antibodies against all VOCs, with markedly enhanced Omicron specific neutralizing antibody responses.

## Introduction

Since late 2019, the Severe Acute Respiratory Syndrome Coronavirus -2 (SARS-CoV-2) virus has led to a global pandemic with over 507 million confirmed infections and 6.2 million deaths, as reported by the World Health Organization (WHO) in April 2022 (1). Despite historic achievements in the distribution of SARS-CoV-2 vaccines, significant gaps remain in the equitable distribution of vaccines with only 15% of people in low income countries having received at least one immunization out of the 11.5 billion doses distributed globally (https://ourworldindata.org/covid-vaccinations). There is also significant concern that booster dosing will also result in significant inequity (2). Combined with the current global dominance of the Omicron VOC (https://www.gisaid.org/phylodynamics/global/nextstrain/, 3) ability to escape humoral immunity (4-6), and the fear of other VOCs yet to emerge due to the pressures of mass vaccination or infection driven immunity, there is a need for the next generation of more broadly protective vaccines to be available in sufficient quantities with superior cold-chain requirements to promote equitable access.

Clover has used Trimer-Tag technology to develop a SARS-CoV-2 vaccine (SCB-2019) with a stabilized prefusion trimeric form of Spike protein (S-Trimer) (7,8). The SCB-2019 vaccine based on the sequence of the Ancestral strain adjuvanted with CpG 1018/Alum has completed clinical phase 1 (NCT04405908) and phase 2/3 SPECTRA trials (NCT04672395). The latter trials enrolled more than 30,000 adult and elderly participants in the Philippines, Colombia, Brazil, South Africa and Belgium, and demonstrated that the SCB-2019 vaccine has a favorable safety and tolerability profile, and significant efficacy against VOCs: 81.7% effective against Delta, 91.8% for Gamma, and 58.6% for Mu against disease of any severity and full protection against severe disease, hospitalization and deaths (9 and 10). An extended follow-up analysis confirms earlier findings and show that SCB-2019 elicited high and durable protection in individuals at approximately six months after the primary vaccination series, including the elderly (Data presented in World Vaccine Congress 2022).

To address Omicron and to drive even broader protection given the potential threat for other VOC to emerge, using the same Trimer-tag platform technology for SCB-2019, we are developing vaccine candidates based on trimerized S-proteins to screen their potential in pre-clinical studies against panels of variants. Based on extensive assessments of immunology and antigenicity, for which the antigenic distance of VOC can be mapped by comparing neutralization values for serum / virus pairs; one can hypothesize that breadth can be achieved by selecting a strain in the centroid range of antigenicity (i.e. the Ancestral strain) and a more distal variant (e.g. Omicron) (11,12). Here we demonstrate that our bivalent vaccine candidate with Spike protein derived from the Ancestral strain (our SCB-2019 vaccine) and the Omicron variant is able to elicit potent cross-protective antibodies against all VOCs, including robust neutralization of Omicron.

## Materials and Methods

### Animal studies, facilities and ethics statements

Specific pathogen-free (SPF) BALB/c female mice (6-8 weeks old) for immunogenicity studies were purchased from Charles River Experimental Animals Co., LTD and kept under standard pathogen-free conditions in the animal care center at Chengdu Hi-tech Incubation Park. All animals were allowed free access to water and diet and provided with a 12 h light/dark cycle (temperature: 16-26°C, humidity: 40% -70%). All mouse experiments were conducted according to international guidelines for animal studies.

### S-Trimer fusion protein expression, purification

S-Trimer fusion proteins SCB-2019 were constructed as previously descripted (14). Similarly, S-Trimer fusion proteins SCB-2022B were constructed utilizing a cDNA encoding the ectodomain of SARS-CoV-2 spike (S) protein from Omicron BA.1 lineage and with a R685A mutation in the furin site, synthesized using Cricetulus griseus (Chinese hamster)-preferred codons by GenScript. The cDNA was subcloned into pTRIMER expression vector (GenHunter Corporation) at *Hin*d *III* and *Bgl II* sites to allow in-frame fusion of the soluble S protein to Trimer-Tag (amino acid residue 1156-1406 from human Type I(α) collagen). The expression vectors were transiently transfected into HEK-293F cell lines (Clover Biopharma) using PEI (Polyscience) and grown in OPM-293 CD05 medium (OPM) with OPM-293 proFeed supplement (OPM). S-Trimer protein was purified to homogeneity from the conditioned medium using Trimer-Tag specific affinity column (Clover Biopharma).

### SEC-HPLC

The purity of S-Trimer was analyzed by Size-Exclusion Chromatography (SEC-HPLC) using Agilent 1260 Infinity HPLC with an analytic TSK gel G3000 SWxL column (Tosoh). Phosphate Buffered Saline (PBS) was used as the mobile phase with OD280 nm detection over a 20 min period at a flow rate of 1 ml/min.

### Receptor binding studies of S-Trimer to human ACE2

The binding affinity of S-Trimer to ACE2 was assessed by Bio-Layer Interferometry measurements on ForteBio Octet QKe (Pall). ACE2-Fc (10 µg/mL) was immobilized on Protein A (ProA) biosensors (Pall). Real-time receptor-binding curves were obtained by applying the sensor in two-fold serial dilutions of S-Trimer (1.125-36 µg/mL in PBS). Kinetic parameters (Kon and Koff) and affinities (KD) were analyzed using Octet software, version 12.0. Dissociation constants (KD) were determined using steady state analysis, assuming a 1:1 binding model for a S-Trimer to ACE2-Fc.

### Vaccine preparation

The test vaccine candidates were formulated with alum (Alhydrogel, Croda, Goole, United Kingdom) plus CpG 1018 (Dynavax Technologies, Emeryville, California). A total of 36 μg of SCB-2019 or SCB-2022B-trimeric protein was mixed first with 900 μg of Alum by gently swirling the mix vial for 30s, then with 1800 μg of CpG 1018, in total 600 μL vol. in vial by gentle inversion 30s at room temperature before administration. Then within 8 hr. 50 μL of vaccine was injected into the hind leg calf muscle per mouse. The bivalent vaccine was prepared with mixture of 18 μg of SCB-2019 and 18 μg of SCB-2022B S-Trimer in 1:1 ratio, then adjuvanted with 900 μg of Alum, inverted gently for 30 seconds and then 1800 μg of CpG 1018 were added, mixed 30 s.

### Animal vaccination

For prime-boost vaccination, Balb/c mice, female (n=10/group) were immunized with SCB-2019, or SCB-2022B 3 μg or Bivalent (1.5 μg of SCB-2019 and 1.5 μg of SCB-2022B) adjuvanted with 75 μg alum plus 150 μg CpG 1018 twice on Day 0 and Day 21. Total 50 μL of vaccine was given each mouse via intramuscular injection. Mice serum was collected on D35.

For three dose boost study, Balb/c mice, female (n=10/group) prime and boost with SCB-2019 3 μg adjuvanted with 75 μg alum plus 150 μg CpG 1018 twice on Day 0 and Day 21, then boosted with 3 μg SCB-2019, or SCB-2022B or Bivalent adjuvanted with 75 μg alum plus 150 μg CpG 1018 on Day 57 via intramuscular injection. Serum was collected on D35 (2 weeks PD2), D56 (Day of 3^rd^ dose boost), D85 (1 month post dose 3), D113 (2 months post dose3) and D141 (3 months post dose 3) for pseudovirus neutralizing antibody test.

### Pseudovirus construction and production

The variants of concern of SARS-CoV-2 spike protein genes were optimized using mammalian codon and synthesized by Genscript, then cloned into pcDNA3.1(+) eukaryotic expression vector. Plasmids encoding Ancestor (Wuhan Hu-1), Alpha (B.1.1.7), Beta (B.1.351), Gamma (P.1), Delta (B.1.617.2), and Omicron (B.1.1.529) SARS-CoV-2 variants S glycoprotein were constructed (mutations compared to the Ancestor were shown in Table 1). The lentiviral packaging plasmid psPAX2 and pLVX-AcGFP-N1-Fluc lentiviral reporter plasmid that expresses GFP and luciferase were obtained from HonorGene (HonorGene, China). Pseudovirions were produced by co-transfection HEK 293T cells with psPAX2, pLVX-AcGFP-N1-Fluc, and plasmids encoding various S genes by using Lipofectamine 3000 (Invitrogen, L3000-015). The supernatants were harvested at 24 ± 2 h post transfection and centrifuged at 1500rpm for 5 min to remove cell debris and then stored at -80°C. Pseudoviruses stock were titrated by infecting 293T-ACE2 cells and luciferase activity was determined following a 44-48 h incubation period at 37°C and 5% CO_2_ by addition Bright-Glo Luciferase Assay System (Promega, E2650) using a microplate reader (TECAN, Spark). Then TCID_50_ of the pseudovirus was calculated according to the Reed-Muench method (13). The virus stock titers were reported in table 1.

**Table 1.** Information of the pseudovirus. Characteristics and information of 7 pseudovirus, including TCID_50_ and mutation on the envelop S protein. Production of variants of concern of SARS-COV-2 virus pseudotype virus stock.

| Table 1: Production of variants of concern of SARS-COV-2 virus pseudotype virus stock. |  |  |  |
| --- | --- | --- | --- |
| PsV Name | Sub-lineage | Stock Titer (TCID<sub>50</sub>/mL) | Mutations On S |
| Ancestor | Wuhan-HU-1 | 1.26x10<sup>5</sup> | / |
| Alpha | B.1.1.7 | 3.79x10<sup>5</sup> | H69-, V70-, Y144-, N501Y, D614G, A570D, P681H, T716I, S982A, D1118H |
| Beta | B.1.351 | 2.19x10<sup>5</sup> | L18F, D80A, D215G, L242-,A243-, L244-, R246I,K417N, E484K, N501Y, D614G, A701V |
| Gamma | P.1 | 2.88x10<sup>5</sup> | L18F, T20N, P26S, D138Y, R190S, K417T, E484K, N501Y, D614G, H655Y, T1027I |
| Delta | B.1.617.2 | 5.85x10<sup>4</sup> | T19R, G142D, E156-, F157-, R158G, L452R, T478K, D614G, P681R, D950N |
| Mu | B.1.621 | 4.21x10<sup>4</sup> | T95I, Y144T, Y145S, Ins146N, R346K, E484K, N501Y, P681H |
| Omicron | B.1.1.529 | 3.79x10<sup>5</sup> | A67V, Δ69-70, T95I, G142D, Δ143-145, Δ211, L212I, ins214EPE, G339D, R346K, S371L, S373P, S375F, K417N, N440K, G446S, S477N, T478K, E484A, Q493R, G496S, Q498R, N501Y, Y505H, T547K, D614G, H655Y, N679K, P681H, N764K, D796Y, N856K, Q954H, N969K, L981F |

### Neutralization assay

Aliquots of test serum samples were first heat-inactivated at 56°C for 30 min, then clarified by centrifugation at 10,000 rcf for 5 min. Samples were serially diluted (3-fold) with assay medium (in 100 µL), incubated with 650 TCID_50_ pseudovirus (in 50 µL) at 37°C for 1 h, along with virus-infected untreated control (virus alone) and cell-alone (background control). Then, freshly-trypsinized 293T-ACE2 cells were added to each well at 20000 cells/well in 100 µL. Following 44-48 h incubation at 37°C in a 5% CO2 incubator, the cells were lysed, and luciferase activity was determined by a Bright-Glo Luciferase Assay System (Promega), according to the manufacturer’s protocol. The IC_50_ neutralizing antibody titer of a given serum sample was defined as the serum dilution where the sample showed the relative light units (RLUs) were reduced by 50% compared to virus-infected control wells. Details of method were reported previously (13).

### Human convalescent serum samples

Human convalescent serum samples from recovered COVID-19 patients were obtained from Public Health Clinical Center of Chengdu in Chengdu, China, under approved guidelines by the Institutional Review Board (IRB), and all patients had provided written informed consent before serum sample were collected. These patients were recently discharged from hospital and the serum was collected at 1-5 weeks after they have been diagnosed as COVID19. Details of sample sourcing and collection are listed in table S1 and certain data previously reported (14).

### Statistical analysis

Data arrangement was performed by Excel and statistical analyses were performed using the Prism 9.2.0 (GraphPad Software). Two-tailed Mann-Whitney tests were used to compare two experiment groups. P values < 0.05 were considered significant. *P < 0.05, **P < 0.01, ***P < 0.001.

## Results

To investigate whether S-Trimer COVID19 vaccine candidates can provide cross-protection against VOCs including Omicron, we have generated a series of SARS-COV-2 pseudoviruses, using Spike protein sequence from the Ancestor (Wuhan-Hu-1), Alpha (B1.1.7), Beta (B.1.351), Gamma (P.1), Delta (B.1.617.2) and Omicron (B.1.1.529) strains (Table 1). Using these pseudoviruses in neutralization assays, we first tested available serum samples collected from convalescent patients. A total of 7 human convalescent sera (HCS) samples (4 moderate, 1 severe, 2 unknown) were initially tested for their pseudovirus neutralizing antibodies against Ancestor, Alpha, Beta, Gamma and Delta; additional human samples (total 35, mild to severe) were accessible later for Ancestor and Omicron only neutralizing antibody testing (Fig. 1). High titers of neutralizing antibodies (IC_50_ GMT over 3 logs) were detected against multiple pseudoviruses, including the Ancestor, Alpha and Gamma strains; neutralizing antibodies against Beta and Delta variants were also maintained at significant levels (IC_50_ GMT 2-3 logs, ∼5-7-fold lower compared to Ancestor). However, neutralization was significantly diminished against Omicron pseudovirus (∼155-fold lower compared to the Ancestor), and only 3 samples out of the total 35 tested were seropositive.

**Fig.1:**
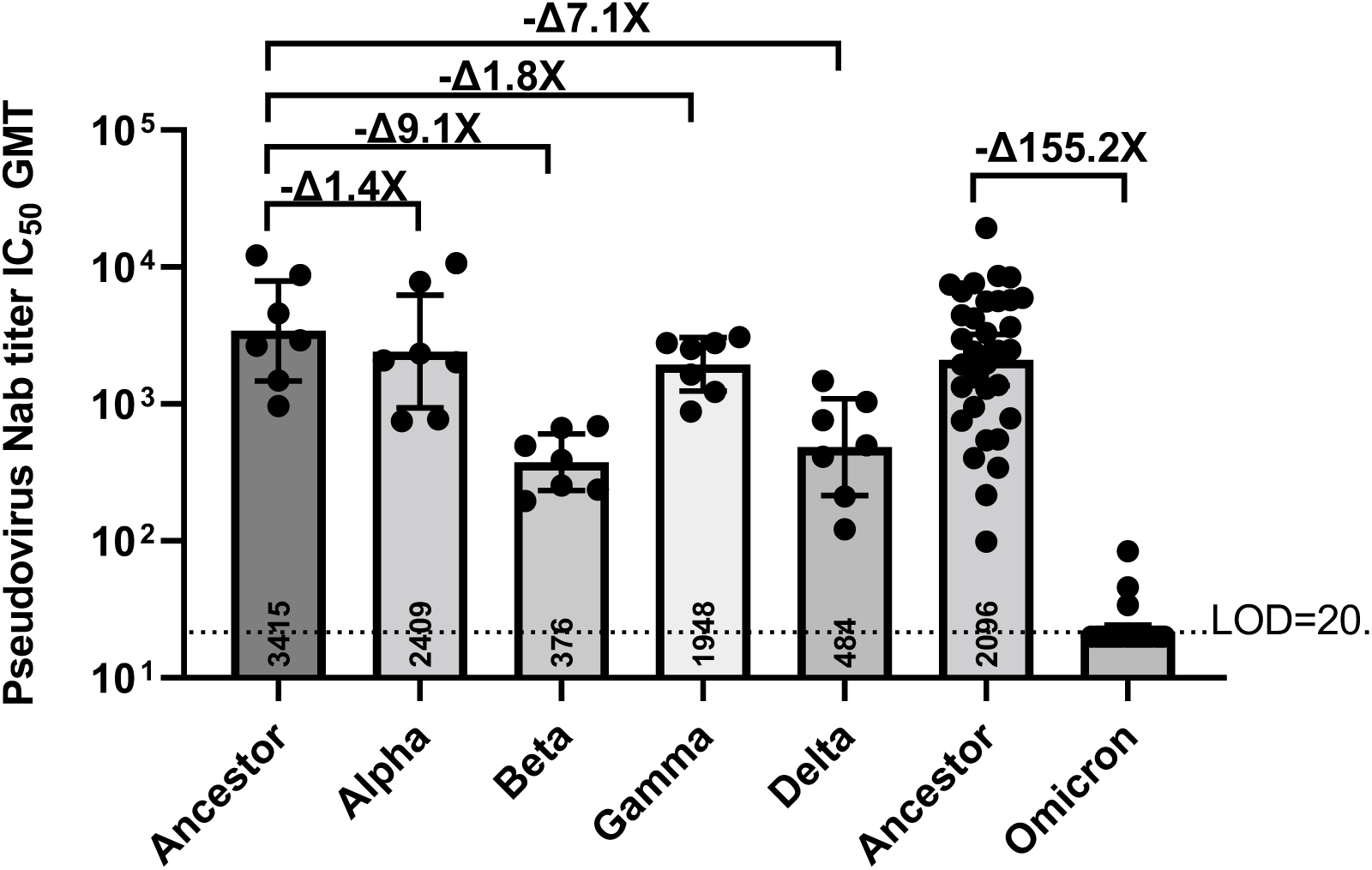
VOCs Pseudovirus Neutralizing antibodies (Nab) titer (IC_50_) from human convalescent sera (HCS). Seven convalescent serum samples (Sample No.1-7 list in table S1) were initially tested against VOCs including Ancestor Hu-1, Alpha, Beta, Gamma and Delta; an additional 35 (Sample No.1-35 list in table S1) serum samples were later tested against Ancestor Hu-1 or Omicron variants. Limit of detection (LOD) titer (IC_50_) is 20. The numbers marked in each bar are the GMT of each test group. Compared with the Ancestral strain, the Nab titer fold decrease for each VOC is labeled as “x”.

To generate a more broadly protective next generation vaccine we first designed SCB-2022B using our Trimer-tag platform based on the Omicron variant full-length Spike protein with an R685A mutation to avoid cleavage at the S1/S2 boundary by furin protease (Fig. 2A). With this mutation, SCB-2022B S-protein produced from CHO cells was intact and showed a clear single band around 250 kDa molecular weight in a reducing SDS-PAGE gel (Fig. 2B) as expected for the trimerized S protein size. The purity was analyzed by size-exclusion SEC-HPLC showing 82% main peak of SCB-2022B S-Trimer respectively (Fig. 2C). The binding affinity (KD) of purified Omicron S-Trimers to the human ACE2 receptor using ForteBio BioLayer interferometry was shown to be 0.8 nM (Fig. 2D). This indicated Omicron S-protein has a high affinity to ACE2 receptor, as previously reported (6). We next generated the bivalent vaccine with a mixture of our SCB-2019 vaccine (14) with the new SCB-2022B S-Trimer in a 1:1 ratio, subsequently formulated with Alum and CpG 1018, the bivalent vaccine contained the same antigen and adjuvant amount compared to the 1^st^ generation vaccine.

**Fig.2:**
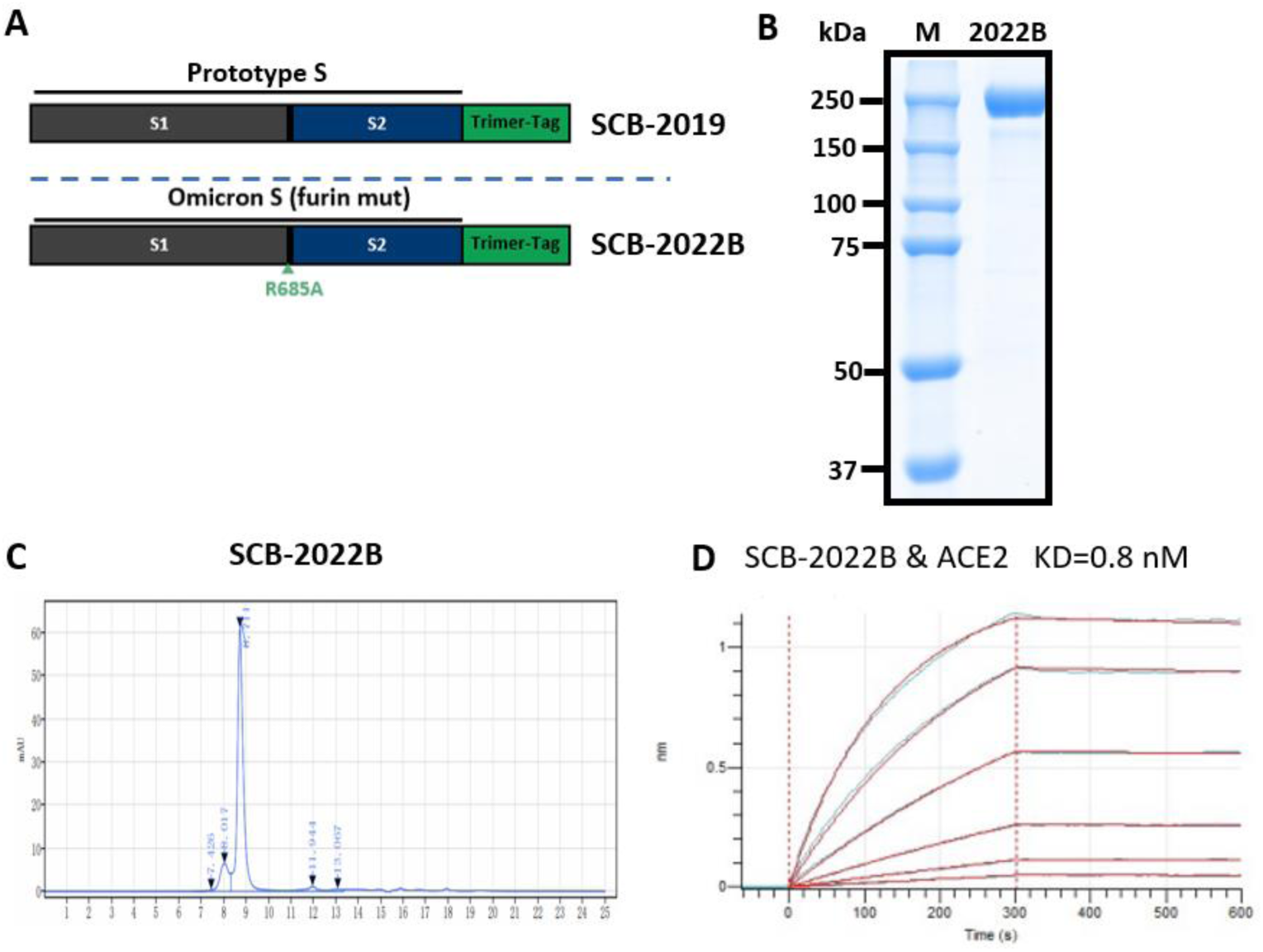
Omicron S-Trimer construction, protein expression and receptor binding affinity. A. Structural design of the trimerized SARS-CoV-2 Omicron spike protein. Schematic representation of the full-length spike protein, SCB-2019: WT ancetor S-Trimer (8). SCB-2022B: a single point mutation R685A at the S1/S2 cleavage site was introduced in the WT Omicron S-Trimer to generate MT S-Trimer. The ectodomain of full-length S is fused with a Trimer-Tag derived from the C-terminal domain of human type I (a) collagen to produce S Trimer. (B) The purified S-Trimer of SCB-2022B was analyzed by Coomassie-stained reducing SDS-PAGE. (C) SEC-HPLC of the purity of Omicron S-Trimer and a small fraction of oligomers and cleaved S1 was shown detached from S-Trimer as indicated. (D)ACE2 receptor binding for SCB-2022B S-Trimer was analyzed by ForteBio BioLayer interferometry indicated.

The immunogenicity of the Omicron monovalent vaccine (SCB-2022B) and bivalent vaccine were then evaluated in a murine two dose prime/boost immunogenicity study and compared with SCB-2019 Ancestor vaccine. Balb/c mice (Female, N=10) were immunized intramuscularly (IM) with total 3 µg of monovalent SCB-2019, or SCB-2022B, or Bivalent constructs all formulated with CpG (150 µg) plus Alum (75 µg). The vaccines were given at study day 0 and 21, serum samples were collected at study day 35 (14 days post-dose 2) and used to determinate the pseudovirus neutralizing antibody responses against VOCs (Fig. 3A). The results indicated two doses of control SCB-2019 Ancestor vaccine can elicit robust neutralizing antibodies against the Ancestor, Alpha, Beta, Gamma and Delta pseudoviruses, but diminished responses against Omicron (Fig. 3B). While SCB-2022B Omicron vaccine immunized mice had significantly higher neutralizing antibodies against Omicron, cross neutralization of other VOCs were lower. However, bivalent vaccine immunized mice had high robust neutralizing antibodies against all VOCs, with significant improvement observed in Omicron specific neutralizing antibodies (about 70-fold increase in GMT), and non-inferiority to others, compared with SCB-2019 even contains only half dose of Ancestor vaccine. This suggests that immunization with the bivalent vaccine can provide enhanced broader protection against VOCs, including the divergent Omicron strain.

**Fig.3:**
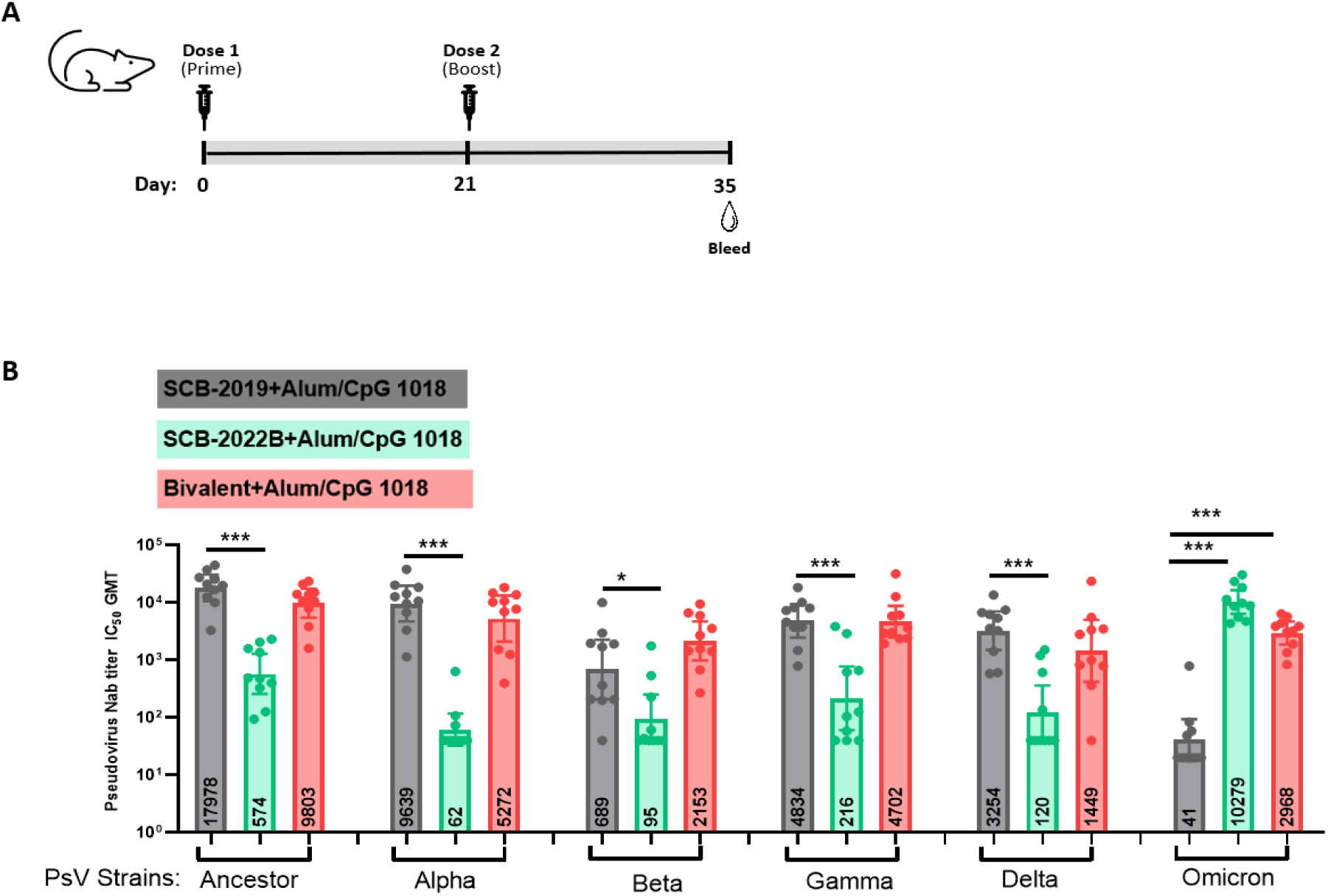
Broadly neutralizing antibody coverage elicited in Bivalent vaccine immunized mice. A. Balb/c mice (n=10/group) were immunized with SCB-2019 (3 μg), SCB-2022B (3 μg) or Bivalent (1.5 μg of SCB-2019 + 1.5 μg of SCB2022B) constructs formulated with 150 μg CpG 1018 plus 75 μg alum twice on Day 0 and Day 21. Serum was collected on D35 for pseudovirus neutralizing antibody testing. B. The study day 35 serum samples were analyzed against VOCs in the pseudovirus neutralization assay (PsVN). Data points represent the pseudovirus neutralizing antibody titer (IC_50_) of the individual animals; Bar horizontal lines indicate geometric mean titers (GMT) for each group ±SEM. The grey bars represent the samples from SCB-2019 immunized mice; Green bars represent the samples from SCB-2022B immunized mice; Red bars represent the samples from SCB-2019 and 2022B bivalent immunized mice. Limit of detection (LOD) titer (IC_50_) is 20. The numbers marked in each bar are the GMT of each test group. For statistical analysis, the comparisons were conducted with Two-tailed Mann– Whitney tests. P values < 0.05 were considered significant. *:P < 0.05, ***:P < 0.001.

Furthermore, to mimic the current situation in humans with many individuals already immunized with ancestor vaccines, and/or infected, we evaluated the immunogenicity of the bivalent vaccine candidate in SCB-2019 pre-immunized animals. Balb/c mice (Female, N=10) were primed and boosted with SCB-2019 formulated with CpG 1018/Alum twice on Day 0 and Day 21, then boosted with SCB-2019, or SCB-2022B or bivalent vaccine formulated with CpG 1018/Alum on Day 57. Serum was collected on D35 (14 days PD2), D56 (day of 3^rd^ boost), D85 (1 month post dose 3, 1MPD3), D113 (2-month post dose3, 2MPD3) and D141 (3-month post dose 3, 3MPD3) for VOCs pseudovirus neutralizing antibody testing (Fig.4A). The results from study day 85 (1MPD3) serum samples indicated, compared with the control group (no 3^rd^ boost), that the 3^rd^ dose boost with the bivalent vaccine significantly enhanced the neutralizing antibody responses against all VOCs, except the Beta variant although such responses were nevertheless robust (Fig. 4B); SCB-2019 monovalent vaccine significantly boosted neutralizing antibodies against Beta, Gamma and Omicron, while the SCB-2022B monovalent vaccine significantly boosted responses against Delta and Omicron.

**Fig.4:**
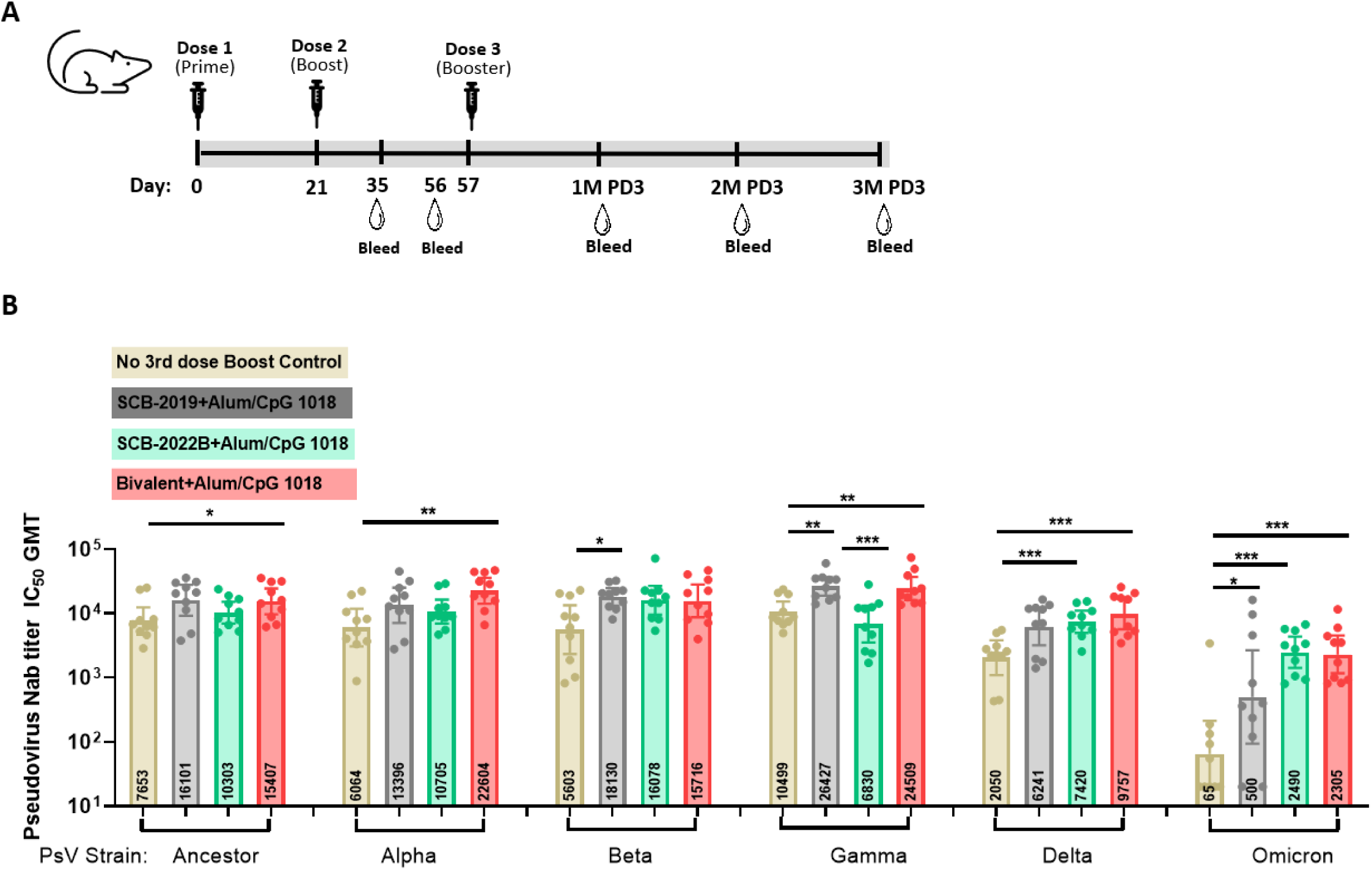
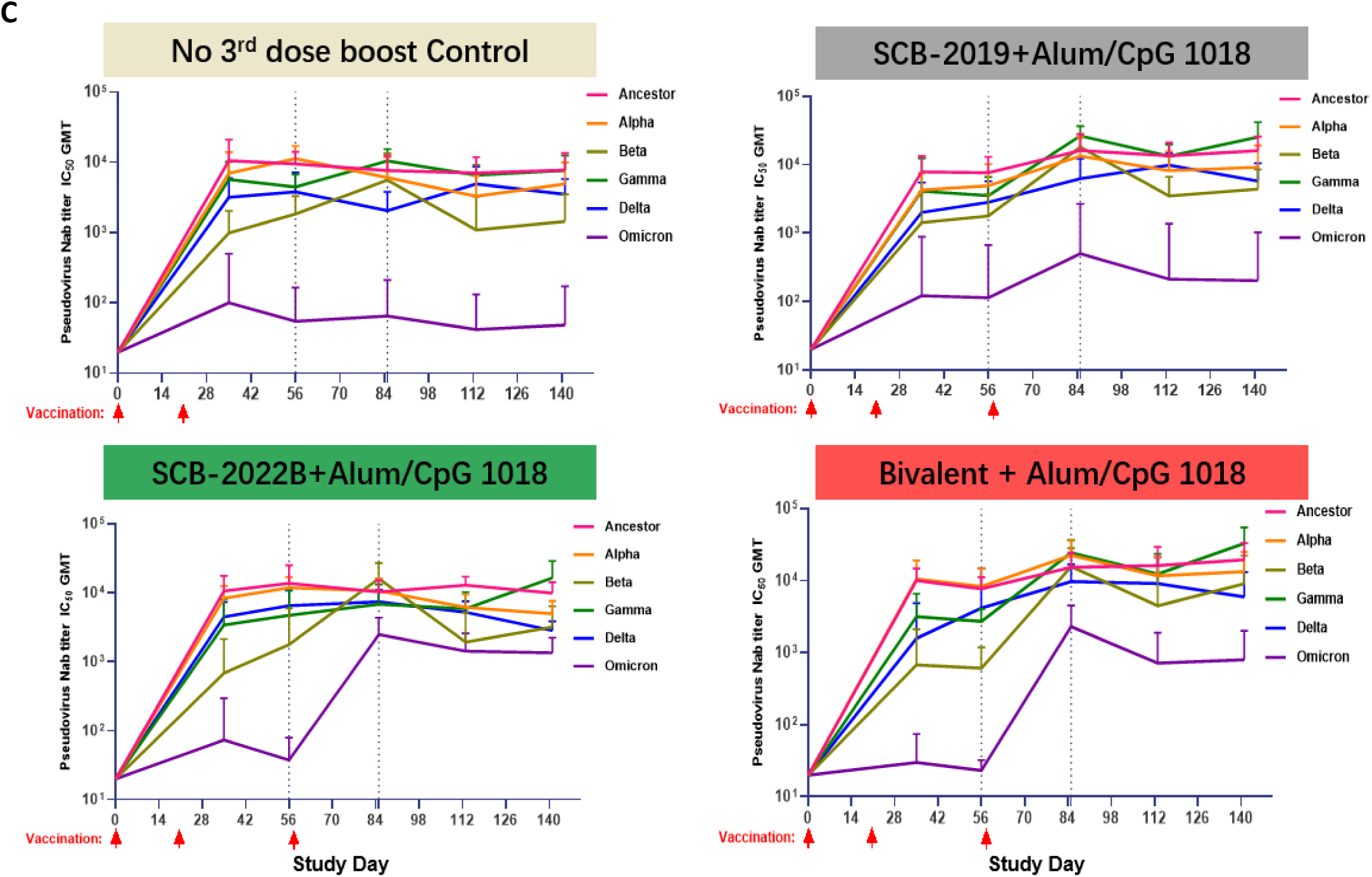
Persistent Broad neutralizing antibody elicited with SCB-2019 (ancestor), SCB-2022B (Omicron) and Bivalent vaccine (SCB-2019+SCB-2022B) as 3^rd^ booster in SCB-2019 prime/boost immunized mice. A. Balb/c mice (n=10/group) were primed and boosted with 3 μg SCB-2019 formulated with 150 μg CpG 1018 plus 75 μg alum on Day 0 and Day 21, then left as control or boosted with SCB2019 (3 μg), SCB-2022B (3 μg) or Bivalent (1.5 μg of SCB-2019 + 1.5 μg of SCB2022B) formulated with 150 μg CpG 1018 plus 75 μg alum twice on Day 57. B. The study day 85 serum samples were analyzed against VOCs via PsV neutralizing assay. Data points represent the pseudovirus neutralizing antibody titer (IC_50_) of the individual animals; Bar horizontal lines indicate geometric mean titers (GMT) for each group ±SEM. Limit of detection (LOD) titer (IC_50_) is 20. The numbers marked in each bar are the GMT of each test group. C. The serum from D0, D35, D56, D85(1M post dose 3), D113 (2M post dose3) and D141 (3M post dose 3) were analyzed with 6 indicated pseudovirus neutralizing for antibody kinetics. The light-yellow bars/box represent the samples from control mice who received no further immunization; Grey bars/box represent the samples from SCB-2019 immunized mice; Green bars/box represent the samples from SCB-2022B immunized mice. Pink bars/box represent the samples from the Bivalent immunized mice. For statistical analysis, the comparisons were conducted with Two-tailed Mann–Whitney tests. P values < 0.05 were considered significant. *:P < 0.05, **:P < 0.01, ***:P < 0.001.

The serum neutralizing responses were monitored post-3^rd^ dose boost over three months to assess the durability of protection (Fig.4C). Serum from the control group (no boost) showed robust neutralizing responses maintained against all VOC except Omicron with a low GMT (95%CI) of 49 (12-1197); SCB-2019 boost significantly improved neutralizing responses against the Ancestral, Alpha, Beta, Gamma and Delta strains, and raised neutralization levels against Omicron, albeit less than the other variants with a GMT (95%CI) of 202 (129-2508) against Omicron. SCB-2022B boost significantly improved neutralizing responses against Omicron with a GMT (95%CI) of 1349 (1324-2112); with a trend for lower set-point responses against other VOCs with comparable GMT titers as the control group (no booster). Bivalent vaccine boost also significantly improved neutralizing responses against all VOC with responses trending higher than the SCB-2022B monovalent boost; with GMT (95% CI) of 799 (762-1973) against Omicron comparable to those elicited by SCB-2022B. These high Omicron specific titers were maintained over the extended observation period (Fig. 4C and Table 2).

**Table 2.** Statistical analysis of the 3^rd^ boost pseudovirus antibody titer.

| PsV Type | Group | Neutralizing antibody titers GMT (95%CI) N=10 |  |  |  |  |  |
| --- | --- | --- | --- | --- | --- | --- | --- |
|  |  | Day 0 | Day 35 | Day 56 | Day 86 | Day 113 | Day 141 |
| Ancestor | Group 1: No 3rd dose boost Control | 20 (/-/) | 10577 (10170-23287) | 9532 (9391-13962) | 7653 (7497-12535) | 7056 (6872-12816) | 7771 (7560-14348) |
|  | Group 2: SCB-2019 CpG 1018/Alum | 20 (/-/) | 7865 (7706-12851) | 7628 (7473-12468) | 16101 (15881-22958) | 13559 (13395-18698) | 15982 (15787-22064) |
|  | Group 3: SCB-2022B CpG 1018/Alum | 20 (/-/) | 10641 (10431-17212) | 13759 (13265-29200) | 10303 (10176-14281) | 12932 (12820-16410) | 9936 (9819-13601) |
|  | Group 4: Bivalent CpG 1018/Alum | 20 (/-/) | 10089 (9984-13368) | 7731 (7607-11621) | 15407 (15188-22248) | 16230 (15752-31169) | 19525 (19209-29412) |
| Alpha | Group 1: No 3rd dose boost Control | 20 (/-/) | 7025 (6749-15664) | 11336 (11223-14886) | 6064 (5923-10488) | 3305 (3242-5291) | 4929 (4792-9196) |
|  | Group 2: SCB-2019 CpG 1018/Alum | 20 (/-/) | 4293 (4222-6495) | 4920 (4800-8663) | 13396 (13136-21507) | 8168 (8046-11979) | 9177 (9009-14405) |
|  | Group 3: SCB-2022B CpG 1018/Alum | 20 (/-/) | 8325 (8196-12338) | 11872 (11702-17170) | 10705 (10537-15958) | 6183 (6078-9469) | 4980 (4894-7684) |
|  | Group 4: Bivalent CpG 1018/Alum | 20 (/-/) | 10541 (10364-16067) | 8323 (8074-16084) | 22604 (22314-31663) | 11756 (11518-19189) | 13356 (13001-24437) |
| Beta | Group 1: No 3rd dose boost Control | 20 (/-/) | 991 (965-1825) | 1838 (1801-2978) | 5603 (5436-10819) | 1097 (994-4297) | 1441 (1333-4830) |
|  | Group 2: SCB-2019 CpG 1018/Alum | 20 (/-/) | 1427 (1391-2543) | 1777 (1746-2747) | 18130 (17972-23089) | 3498 (3431-5593) | 4416 (4316-7520) |
|  | Group 3: SCB-2022B CpG 1018/Alum | 20 (/-/) | 686 (647-1913) | 1781 (1701-4268) | 16078 (15700-27906) | 1926 (1818-5306) | 3190 (3096-6134) |
|  | Group 4: Bivalent CpG 1018/Alum | 20 (/-/) | 676 (639-1840) | 608 (597-971) | 15716 (15418-25050) | 4482 (4142-15086) | 9113 (8688-22403) |
| Gamma | Group 1: No 3rd dose boost Control | 20 (/-/) | 5692 (5583-9123) | 4474 (4423-6070) | 10499 (10363-14731) | 6585 (6517-8705) | 7699 (7555-12186) |
|  | Group 2: SCB-2019 CpG 1018/Alum | 20 (/-/) | 4125 (4022-7372) | 3534 (3477-5342) | 26427 (26150-35086) | 13288 (13140-17895) | 25355 (24863-40755) |
|  | Group 3: SCB-2022B CpG 1018/Alum | 20 (/-/) | 3414 (3270-7914) | 4722 (4501-11630) | 6830 (6669-11845) | 5903 (5779-9773) | 16461 (16103-27663) |
|  | Group 4: Bivalent CpG 1018/Alum | 20 (/-/) | 3165 (3083-5738) | 2727 (2698-3661) | 24509 (24116-36787) | 12403 (12005-24844) | 33037 (32450-51404) |
| **Delta** | Group 1: No 3rd dose boost Control | 20 (/-/) | 3196 (3113-5800) | 3824 (3768-5554) | 2050 (2018-3066) | 4914 (4840-7237) | 3521 (3473-5027) |
|  | Group 2: SCB-2019 CpG 1018/Alum | 20 (/-/) | 2009 (1957-3648) | 2832 (2787-4219) | 6241 (6133-9618) | 9879 (9787-12759) | 5768 (5694-8085) |
|  | Group 3: SCB-2022B CpG 1018/Alum | 20 (/-/) | 4482 (4406-6861) | 6537 (6422-10127) | 7420 (7335-10074) | 5244 (5188-6977) | 2839 (2810-3754) |
|  | Group 4: Bivalent CpG 1018/Alum | 20 (/-/) | 1581 (1518-3547) | 4195 (4070-8119) | 9757 (9594-14854) | 9162 (8852-18854) | 5961 (5749-12589) |
| **Omicron** | Group 1: No 3rd dose boost Control | 20 (/-/) | 101 (58-1440) | 55 (39-547) | 65 (44-727) | 42 (20-740) | 49 (12-1197) |
|  | Group 2: SCB-2019 CpG 1018/Alum | 20 (/-/) | 122 (-47-5392) | 114 (51-2083) | 500 (390-3929) | 213 (123-3023) | 202 (129-2508) |
|  | Group 3: SCB-2022B CpG 1018/Alum | 20 (/-/) | 73 (42-1049) | 37 (34-147) | 2490 (2448-3815) | 1425 (1402-2134) | 1349 (1324-2112) |
|  | Group 4: Bivalent CpG 1018/Alum | 20 (/-/) | 30 (23-245) | 23 (23-36) | 2305 (2237-4403) | 716 (683-1751) | 799 (762-1973) |

## Discussion

In this study, we corroborated other reports (15-18) that human convalescent sera have substantially lower levels of Omicron neutralizing antibodies compared to Ancestral strain, although the same sera generally maintain broadly cross-reactive neutralizing antibodies against other VOCs. This verified the utility of our panel of VOCs in our neutralization assay to assess the consequences of the Omicron S-protein mutations on humoral immunity (19). The evidence of breakthrough infections in fully vaccinated individuals further emphasizes the importance of booster doses and potential of next generation vaccines to enhance protection against divergent VOCs such as the Omicron lineage (4,5).

Therefore, we designed a bivalent vaccine, to address advanced models that attempt to define the antigenic range of SARS-CoV-2 (20) by selecting our Ancestor vaccine, SCB-2019, which occupies a centroid range of antigenicity and has shown high levels of efficacy against VOCs in a phase 3 trial (10), and the Omicron strain given its divergent antigenicity and its global prevalence. This approach is supported by the observation that individuals who have received two doses of Ancestor vaccine and subsequently infected with Omicron have high levels of broad cross-reactive neutralization against panels of VOCs (21). An Omicron only monovalent vaccine, while likely to elicit protection against Omicron and related sub-lineages, is however potentially ineffective against other VOC distal from its antigenic position; a rationale for bivalency to mitigate this risk. The subsequently derived Omicron component of the bivalent approach, SCB-2022B, using the same Trimer-Tag technology as SCB-2019, appears trimeric in nature and binds with high affinity to the ACE-2 receptor. Combined with SCB-2019 at a 1:1 ratio to create the bivalent vaccine, formulated with CpG/alum. The bivalent vaccine keeps the total same amount of antigen and adjuvant as the 1^st^ generation vaccine, therefore not impact the supply. The murine preclinical studies allowed us to assess the breadth of cross neutralization of a panel of VOCs including Omicron.

In a priming two-dose schedule setting, even only contains half dose of Ancestor variant vaccine, the bivalent vaccine was able to elicit robust cross neutralization of all VOCs (range 10^3^-10^4^ titers, non-inferior to those elicited by monovalent Ancestor vaccine), including high titers against Omicron (IC_50_ GMT 2968) in the mice. In animals primed with our SCB-2019 vaccine, the bivalent vaccine was able to elicit strong booster neutralization responses against all VOCs tested with substantial levels against Omicron (∼10^3^ titer range) that were sustained up to 3 months post boosting. The SCB-2019 and SCB-2022B monovalent formulations also boost neutralization robustly, the latter as expected particularly well against Omicron. In totality, the bivalent vaccine trended towards incrementally superior titers against the panel of VOCs during the evaluation of humoral kinetics post boosting; however, the third immunization with SCB-2022B and the bivalent vaccine both elicited Omicron neutralization responses with peak and set-point titers below the other VOCs. This is suggestive of the original antigenic sin hypothesis, in which adaptive immunity is partially imprinted against the initial antigens presented to the naïve immune system (22).

In conclusion, despite effectiveness data demonstrating that ancestor vaccines remain highly effective against severe disease and hospitalization caused by Omicron, there is an opportunity to improve protection against all cause disease and transmission caused by the currently dominant Omicron strain. Our murine preclinical priming and booster studies demonstrate the value of a bivalent formulation, combining our SCB-2019 vaccine with an Omicron specific SCB-2022B construct, to elicit broad neutralization coverage against a panel of VOCs including Omicron. Given gaps that remain in equitable vaccination in low-income countries, the ongoing major outbreaks in China and the threat of future VOCs, a broadly protective bivalent vaccine with the clinical tolerability and thermal stability profile of our SCB-2019 vaccine, would contribute towards public health goals.

## Acknowledgement

The authors would like to acknowledge and thank Public Health Clinical Center of Chengdu for supply the human convalescent sera samples.

## Author Contribution

J.G.L, P.L. N.J. and R.X conceived this project, and R.X and D.S. designed the study. D.S. oversaw mouse studies, cell culture for antigen production and developed in vitro antibody/neutralizing antibody assays. X.L and C.H. performed expression vector construction and antibody titer experiments. C.Z conducted protein purification experiments. X.H and Z.M performed antigen production. Q.W. and W.Q. performed the animal studies.

---

**Table S1.** Human Convalescent Sera(HCS) Sample Information.

| Table S1: Human Convalescent Sera (HCS) Sample Information |  |  |  |  |  |  |  |
| --- | --- | --- | --- | --- | --- | --- | --- |
| Sample No. | Patient ID. | Sample collection Time | Patient Age | Gender | Mild/Moderate/Severe | Admission Date | Signed ICF or not |
| 1 | 38 | 03/11/2020 | 37 | M | Severe | 01/29/2020 | Yes |
| 2 | 56 | 03/11/2020 | 55 | M | Moderate | 02/08/2020 | Yes |
| 3 | K57 | 03/04/2020 | 54 | F | Moderate | 02/08/2020 | Yes |
| 4 | K62 | Unknown | Unknown | Unknown | Unknown | Unknown | Yes |
| 5 | K85 | Unknown | Unknown | Unknown | Unknown | Unknown | Yes |
| 6 | K88 | 03/04/2020 | 47 | M | Moderate | 02/12/2020 | Yes |
| 7 | 110 | 03/15/2020 | 23 | M | Moderate | 02/22/2020 | Yes |
| 8 | 94s | 02/16/2020 | 55 | F | Moderate | 02/12/2020 | Yes |
| 9 | 1s | 02/06/2020 | 63 | F | Mild | 01/23/2020 | Yes |
| 10 | 10s | 02/05/2020 | 47 | M | Severe | 01/26/2020 | Yes |
| 11 | 21s | Unknown | 34 | M | Mild | 01/25/2020 | Yes |
| 12 | 31s | unknown | 65 | M | Mild | 01/30/2020 | Yes |
| 13 | 36s | 02/05/2020 | 62 | F | Mild | 01/29/2020 | Yes |
| 14 | 37s | Unknown | 67 | F | Mild | 01/29/2020 | Yes |
| 15 | 39s | Unknown | 25 | M | Mild | 01/29/2020 | Yes |
| 16 | 60s | 02/15/2020 | 41 | M | Mild | 02/08/2020 | Yes |
| 17 | 61s | 02/15/2020 | 27 | F | Mild | 02/09/2020 | Yes |
| 18 | 93s | 02/15/2020 | 62 | F | Moderate | 02/12/2020 | Yes |
| 19 | 95s | 02/17/2020 | 35 | M | Moderate | 02/12/2020 | Yes |
| 20 | 3 | 03/11/2020 | 60 | M | Severe | 01/26/2020 | Yes |
| 21 | 81 | 03/11/2020 | 59 | M | Moderate | 02/12/2020 | Yes |
| 22 | 83 | 03/15/2020 | 50 | M | Moderate | 02/12/2020 | Yes |
| 23 | 108 | 03/11/2020 | 57 | F | Moderate | 02/21/2020 | Yes |
| 24 | 109 | Unknown | 36 | F | Moderate | 02/21/2020 | Yes |
| 25 | 117 | 03/15/2020 | 74 | F | Moderate | 02/28/2020 | Yes |
| 26 | 130 | 03/15/2020 | 47 | F | Moderate | 03/05/2020 | Yes |
| 27 | K27 | 03/04/2020 | 66 | M | Mild | 01/25/2020 | Yes |
| 28 | K80 | unknow | unknown | unknown | unknown | unknown | Yes |
| 29 | K87 | 03/04/2020 | Unknown | F | Moderate | 02/12/2020 | Yes |
| 30 | K92 | 03/04/2020 | 59 | F | Moderate | 02/12/2020 | Yes |
| 31 | K111 | Unknown | Unknown | Unknown | Unknown | Unknown | Yes |
| 32 | K114 | Unknown | Unknown | Unknown | Unknown | Unknown | Yes |
| 33 | K120 | 03/04/2020 | Unknown | F | Mild | 02/29/2020 | Yes |
| 34 | K121 | Unknown | Unknown | Unknown | Unknown | Unknown | Yes |
| 35 | K122 | Unknown | Unknown | Unknown | Unknown | Unknown | Yes |

